# Probing mechanotransduction in living cells by optical tweezers and FRET-based molecular force microscopy

**DOI:** 10.1101/2021.01.24.427943

**Authors:** M. Sergides, L. Perego, T. Galgani, C. Arbore, F.S. Pavone, M Capitanio

**Author notes:** Correspondence and requests for materials should be addressed to M.C.

## Abstract

Cells sense mechanical signals and forces to probe the external environment and adapt to tissue morphogenesis, external mechanical stresses, and a wide range of diverse mechanical cues. Here, we propose a combination of optical tools to manipulate single cells and measure the propagation of mechanical and biochemical signals inside them. Optical tweezers are used to trap microbeads that are used as handles to manipulate the cell plasma membrane; genetically encoded FRET-based force sensors inserted in F-actin and alpha-actinin are used to measure the propagation of mechanical signals to the cell cytoskeleton; while fluorescence microscopy with single molecule sensitivity can be used with a huge array of biochemical and genetic sensors. We describe the details of the setup implementation, the calibration of the basic components and preliminary characterization of actin and alpha-actinin FRET-based force sensors.

## 1 Introduction

The study of the mechanical regulation of biological systems has greatly expanded in the last decade. There is a large body of evidence indicating that the mechanical properties of the extracellular environment directly affect the mechanical properties of the cell and, in turn, activate signalling pathways that switch specific genes and feedback programs on or off. A well-established example in this respect is the YAP (Yes-associated protein) and TAZ (transcriptional co-activator with PDZ-binding motif) complex, whose transcriptional activity and cell localization have been shown to be strictly related to the mechanical properties of the extracellular environment[1]. In fact, it has been shown that when various cell types are cultured on soft substrates, YAP–TAZ localizes in the cytoplasm and its transcriptional activity is low and comparable to YAP–TAZ knockdowns. On the other hand, when cells are cultured on stiff substrates, YAP–TAZ moves into the nucleus and its transcriptional activity is greatly enhanced and similar to that of cells grown on plastic[1].

One important route for cell mechanical sensing occurs through adhesion complexes, which bind to the extracellular matrix (through focal adhesions) and/or to neighbour cells in a tissue (through adherens or tight junctions). Although the membrane proteins involved in the different adhesion complexes vary greatly both in structure and function (integrins, cadherins, and ZO-1 for focal adhesions, adherens junctions and tight junctions, respectively), all these complexes exhibit a dynamic interaction of their cytoplasmic domain with the actin cytoskeleton. This interaction is mediated by numerous adaptor proteins that create a bridge between the adhesion molecules and actin. The dynamic nature of this link is fundamental in regulating the strength of cell adhesions to adapt to the changes in the mechanics of the environment and during the different phases of cell’s life [2]. For example, it has been recently shown that, during tissue growth, a fluid-to-solid transition (usually named unjammed-to-jammed transition) occur in cultured epithelial monolayers [3] and *in-vivo* [4]. Jamming in epithelial monolayers occurs at a critical cell density and it is associated with changes in cell shape and mechanical forces between adjacent cells. Cell unjamming has been shown to be associated with RAB5A-induced reawakening of collective cell motility [5]. Jamming-to-unjamming transitions are also critical in tumour progression and cancer invasion. In cancer, tumour tissue is generally stiffer than the surrounding healthy tissue, however tumour cells are softer than healthy cells. The mechanical properties of cancer cells and their extracellular matrix have been shown to play a critical role in unjamming transitions and metastasis[6]–[8]. The complexity of these mechanisms and the involvement of different mechanical, biochemical, and genetic signals, makes its study extremely complex and virtually impossible by using a single experimental technique or approach.

Therefore, it has become increasingly important to develop and combine advanced experimental methodologies and setups to uncover how mechanical signals are sensed by cells, how they propagate inside the cell and to the cell’s cytoskeleton, and how they eventually lead to changes in biochemical signalling and gene expression. To this end, we have implemented an all-optical experimental apparatus that combines manipulation of the cell, imaging of intracellular tension on specific proteins, and detection and tracking of biochemical and genetic signals with single molecule sensitivity.

Optical manipulation is obtained by using optical tweezers. Optical tweezers are made of a laser beam tightly focussed by the microscope objective[9]–[11]. Gradient forces acting near the laser focus attract micrometer-sized particles that can be used as handles to manipulate single molecules[12] and/or cells[13]. Optical tweezers are a versatile tool that allows manipulation as well as measurement of forces, which can be applied from both the outer plasma membrane or inside the cell. By coating the trapped microsphere with adhesion molecules such as fibronectin or cadherin, focal adhesions or adherens junctions, respectively, can be specifically stimulated on the plasma membrane with mechanical forces of different intensity and frequency (Fig.1). The relative ease of applying and measuring physiologically relevant forces have made optical tweezers one of the ideal tools to study cell mechanics.

**Figure 1:**
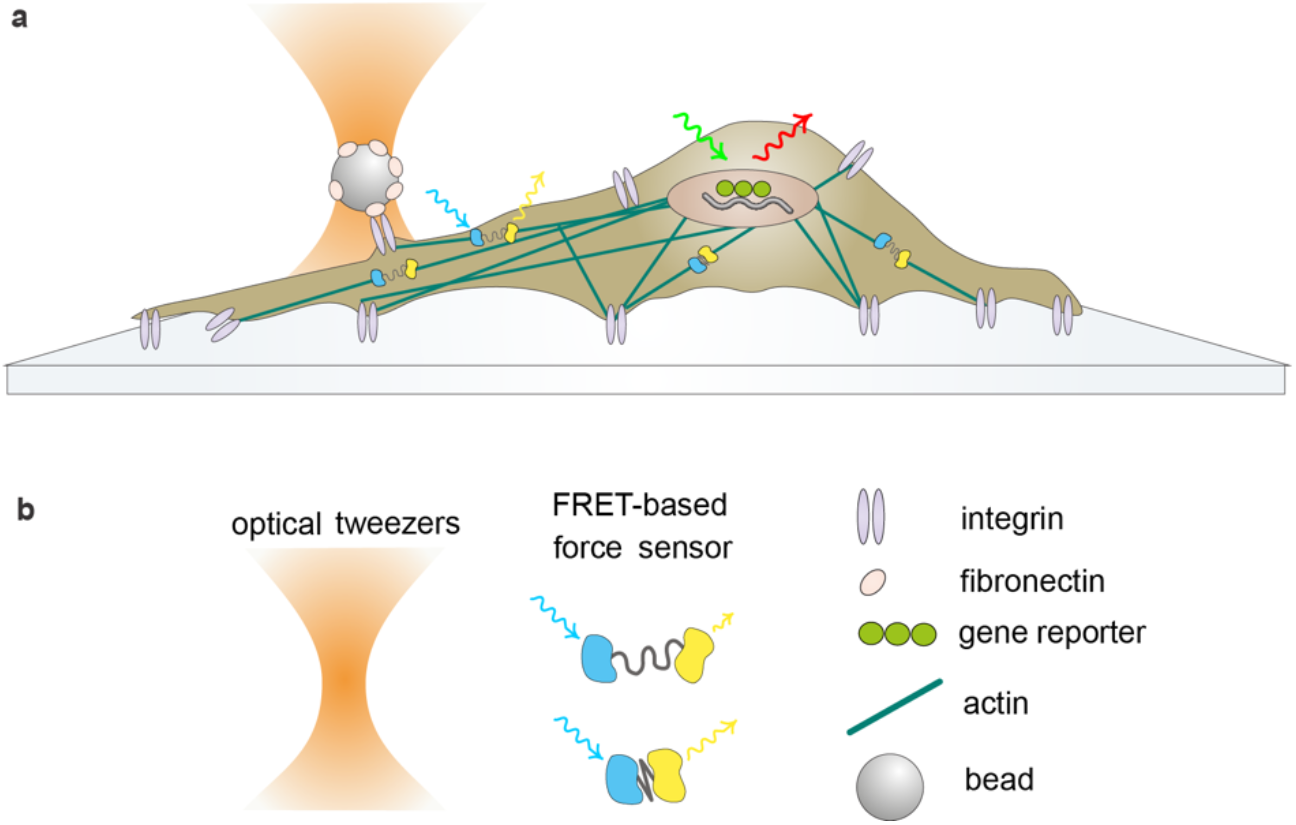
Schematic of the experimental approach used in mechanotransduction experiments. **(a)** Optical tweezers are used to apply and measure mechanical load on specific adhesion molecules on the cell plasma membrane. Genetically encoded FRET-based force sensors measure the propagation of mechanical signals in the cell. Fluorescent gene reporters detect mechanically-induced changes in gene expression. **(b)** The various components used in a mechanotransduction experiment

On the other hand, the transmission of mechanical forces within the cell can be probed by using genetically encoded FRET-based force sensors. FRET-based force sensor modules are usually made by a donor-acceptor fluorescent protein pair linked together by an elastic peptide[14]–[16]. The DNA sequence coding for the force sensor module is inserted within the coding sequence of the protein of interest and transfected in living cells. When the recombinant protein is expressed by the cell and set under tension, the elastic peptide extends under force, the distance between the donor and the acceptor increases, and the FRET efficiency consequently decreases, allowing the measurement of tension changes in different regions of the cell and/or under different conditions of mechanical stimuli (Fig. 1). Force sensors for several proteins (among which actin[17], spectrin[18], alpha-actinin[18], alpha-catenin[19], nesprin[20], vinculin[21]) and force sensors modules based on different linkers have been developed[18], [21]–[23], allowing the measurement of different ranges of forces from few pN to several tens of pN per molecule.

Eventually, mechanical signals are converted into biochemical and genetic signals. Our setup is designed to allow detection of fluorescent signals with single molecule sensitivity. This capability can be exploited to detect several kinds of biochemical signals triggered by mechanical cues (i.e. ion fluxes such as Ca2+ or nuclear shuffling of transcription factors such as YAP/TAZ). Alternatively, fluorescent reporters of gene expression can be used to monitor changes induced by mechanical stimuli in real time, both for protein[24] and mRNA[25]–[27] expression, with sensitivity up to the single molecule (Fig.1).

Here, we describe the details of the setup implementation and how to efficiently combine the different techniques; the calibration of the basic components; as well as preliminary characterization of cell manipulation by optical tweezers, intracellular tension measurement using actin and alpha-actinin FRET-based force sensors, and detection of chromophores with single molecule sensitivity.

## 2 Experimental Setup Design

In order to directly observe and image the response of biological systems to mechanical stimulation, the experimental arrangement combined optical tweezers and fluorescence microscopy. A schematic of the apparatus is shown in Fig. 2. The system allowed for the simultaneous manipulation of specific cell membrane receptors and detection of intracellular signals, with sensitivity down to the single molecule. The setup was realized around a commercial infinity corrected microscope (Nikon Eclipse Ti-U) which could be adjusted to accommodate different excitation and emission wavelengths as well as highly inclined laminated optical sheet (HILO) and total internal reflection fluorescence (TIRF) microscopy techniques through custom components mounted on an optical table[28]. The microscope objective (Nikon CFI Apochromat TIRF, 60x magnification and numerical aperture *NA* = 1.49), provided high transmission and efficient correction of chromatic aberrations in the range from 435 nm to 1064 nm, which enabled optimal combination of the imaging of single molecules in the visible spectrum and trapping particles with a near-infrared laser (808 nm wavelength) on the same focal plane. The microscope was equipped with a piezoelectric stage (PI PInano P-545) with the capability to translate the sample along the three spatial directions (200 μm x 200 μm x 200 μm) with sub-nanometric resolution and step times of the order of a millisecond, which enabled precise positioning and nanometer stabilization of the sample [29]. In addition, the piezoelectric stage was mounted on top of a high stable stage (PI M-545), capable of long range displacements of 25 mm x 25 mm. The optical trap could be moved independently from the sample stage by steering the laser source using a piezoelectric mirror, thus allowing the decoupling of fluorescence imaging from optical trapping. Furthermore, the nanometric trap positioning combined with three-dimensional position detection using a quadrant detector photodiode (QPD), allowed for the precise application and measurement of external forces on cell membranes. The movement of the stage, the operation of the laser sources and collection of data via the various detectors could be controlled remotely by a custom-made computer software. For data acquisition, an acquisition board (National Instruments PCIe-6343) was used, consisting of 32 analogue inputs with 16-bit resolution and sampling rate of 500 kS/s, as well as 4 analogue outputs with the same resolution and sampling rate of 900 kS/s. The optical setup was positioned on an anti-vibration optical table (Melles Griot) to minimize mechanical vibrations.

**Figure 2:**
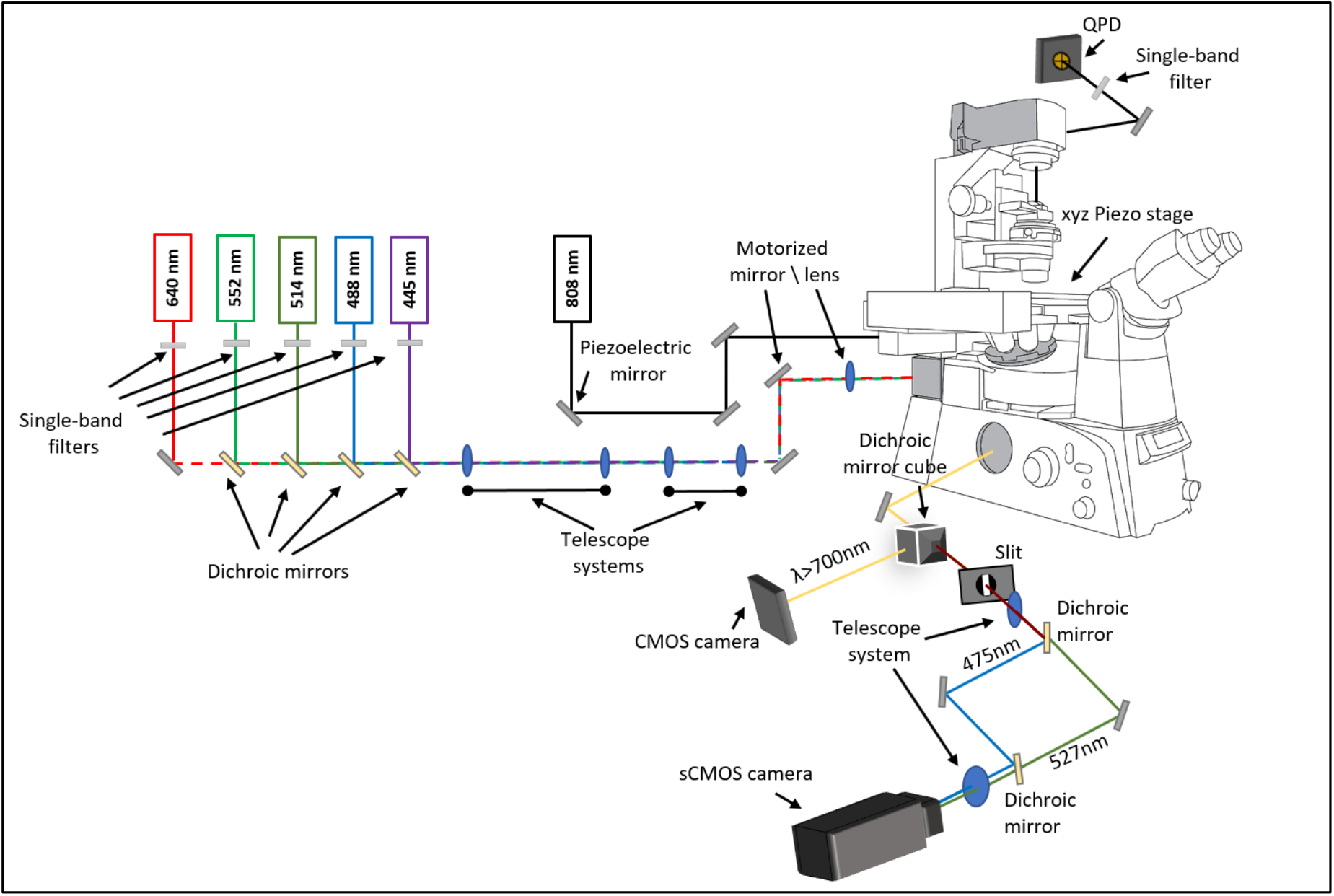
Schematic of the experimental setup used in mechanotransduction experiments. The apparatus combined dual-colour fluorescence microscopy and optical manipulation by optical tweezers.

### 2.1 Fluorescence Microscopy

The fluorescence microscopy part of the experimental apparatus consisted of five excitation semiconductor TEM00 laser sources (Fig. 3a). In particular, from right to left, an OBIS 445LX laser at 448 nm, with a maximum output power *P*_*max*_ = 75 mW and a collimated beam waist *ω* = (0.6 ± 0.1) mm, an OBIS 488LX at 488 nm, with *P*_*max*_ = 150 mW and *ω* = (0.7 ± 0.1) mm, an OBIS 514LX at 518 nm, with *P*_*max*_ = 40 mW and *ω* = (0.6 ± 0.1) mm, an OBIS 552LS at 552 nm, with *P*_*max*_ = 150 mW and *ω* = (0.70 ± 0.05) mm, and finally an OBIS 640LX laser at 643 nm with *P*_*max*_ = 150 mW and *ω* = (0.8 ± 0.1) mm. Telescope systems were placed after each laser source output, in order to enlarge the laser beams to a common size and optimize their spatial profile through a pinhole placed in the focal plane between the two lenses of the telescope. Narrow band-pass filters (Semrock FF01-445/20-25, FF01-488/6-20, FF01-513/13-25, FF01-542/27-25, FF01-640/14 respectively) were also positioned at the output of all diode laser sources to eliminate undesired wavelengths that exist due to the spurious emission outside the emission bandwidth. It is noted that in the absence of the filters, the spurious emissions constituted an important source of noise in single molecule fluorescence microscopy experiments. Each beam was directed to the microscope by dichroic mirrors (Semrock Di02-R442, LM01-503-25, Di02-R514 and Chroma ZT568rdc) allowing for the consecutive excitation of samples by the different wavelengths. Moreover, two achromatic telescope systems were introduced. The two telescopes enlarged the excitation beams to allow for full-field (200 μm x 200 μm) excitation at low powers, suitable for cell imaging. The second telescope system was placed on flip mounts to provide the option for half-field view at 4x power when removed, optimal for single molecule imaging. An additional achromatic lens focused the excitation beam into the back aperture of the objective lens. This achieved constant intensity illumination since the focussing lens and the objective were in a telescope configuration. An iris was placed on the back-focal plane of the focussing lens and, thus, conjugated to the objective’s focal plane to spatially limit the excitation beam and regulate the field of illumination. By placing a mirror before the focussing lens, and the focussing lens itself on motorized translators, it was possible to translate the excitation beam at the entrance of the objective to switch between different illumination schemes such as wide field, HILO and TIRF.

**Figure 3:**
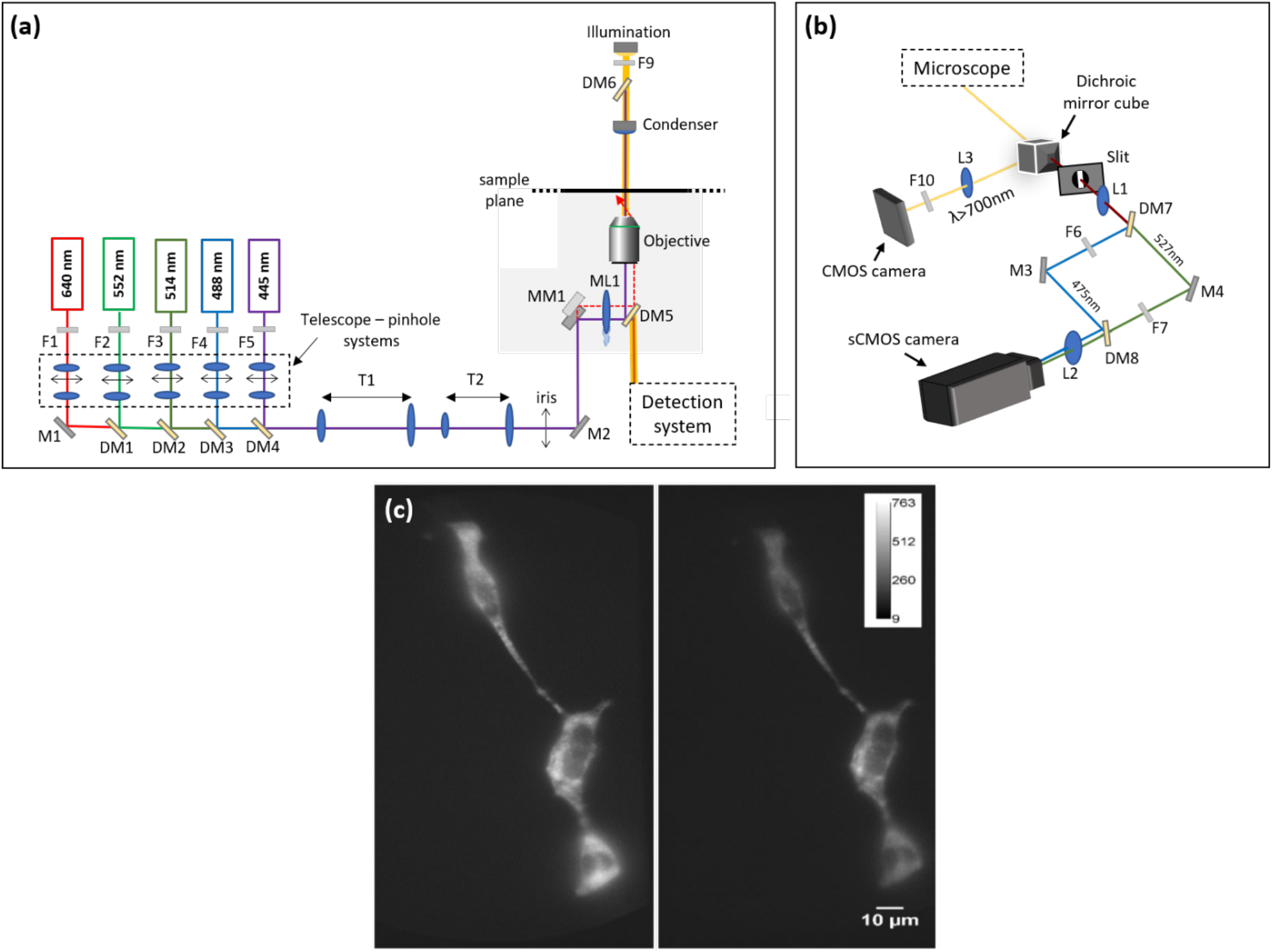
**(a)** Schematic of the fluorescence excitation part of the experimental apparatus containing five laser sources. After each laser system narrow band filters F1 – F5 were placed to eliminate undesired wavelengths. Telescope/pinhole systems were also placed to enlarge the excitation beams to a common size and optimize their spatial profile. With the use of dichroic mirrors DM1-DM4 the excitation beams were directed to the microscope after passing through two telescope systems, T1 and T2, which allowed for different fields of excitation. Motorized mounts were used for mirror MM1 and focusing lens ML1 providing the possibility to switch between different illumination schemes (wide field, HILO and TIRF). The dichroic mirror DM5 could be selected as such with the aid of the microscope wheel to match the excitation and emission wavelengths necessary. Filter F9 was used to allow wavelengths longer than 700 nm to reach the sample, making possible bright field imaging via the detection system. **(b)** The detection part of the system allowed for simultaneous imaging of the bright field and fluorescence emission. The light exiting the microscope setup was split into two parts by a dichroic mirror cube. Wavelengths longer than 700 nm were reflected on a CMOS camera for bright-field imaging. Shorter wavelengths passed through a slit in order to split the fluorescence image to two parts. Dichroic mirror DM7 transmitted wavelengths longer than 506 nm and reflected shorter ones. The two paths were then recombined by DM8 identical to DM7 and focused to the sCMOS camera side by side by the achromatic doublet telescope system consisting lenses L1 and L2. Emission filters F6 and F7 were placed in each arm to filter out wavelengths other than desired emission signal. **(c)** Live HEK cells transfected by the cpstFRET sensor imaged in two emission channels simultaneously. On the left, the “acceptor” channel is shown corresponding to 542 nm emission and on the right the “donor” channel corresponding to 479 nm emission.

The commercial microscope contained a microscope wheel which could accommodate several dichroic mirrors (DM5) (e.g. Chroma ZT440/488/561/635rpc-uf2, Chroma ZT488/561rpc-uf2, Chroma ZT440/514/561/640rpc-uf1) reflecting the excitation beams to the objective, as well as transmitting the emitted fluorescence to the detection system. This made it easy to adapt to different excitation wavelengths for samples characterized by the presence of a combination of fluorophores. The detection apparatus was able to image two emission channels of the fluorophores on a single sCMOS camera (Hamamatsu Orca-Flash4.0 V2) (Fig. 3b). This was particularly beneficial for the realization of FRET experiments since it was essential to concurrently display the emission from both donor and acceptor fluorophores. Simultaneous bright field imaging was achieved by a CMOS camera (Thorlabs DCC1545M). This can be particularly useful to perform combined trapping and fluorescence experiments. In order to achieve this, the halogen illumination was filtered with a long-pass filter (Thorlabs FEL0700) to allow the transmission of wavelengths longer than 700 nm to minimize interfere with fluorescence microscopy. The light exiting the microscope arrived at a dichroic mirror (Chroma ZT775sp-2p), which was transparent to wavelengths shorter than 730 nm and reflective for longer wavelengths. The reflected light was focused on the CMOS camera after passing through a notch filter (Thorlabs NF808-34) to block the optical tweezers beam. The transmitted light passed through the centre of a slit positioned on the image plane produced by the microscope tube lens. This was done to divide the image into two identical halves centred side-by-side on the sCMOS camera chip. Following the slit, a 1:1 telescope system was devised consisting of two achromatic doublet lenses with the same focal length. The first lens of this system was placed at a distance equal to its focal length from the slit while the second one was placed at the same distance from the sCMOS camera. A dichroic mirror (Semrock FF506-Di03) reflected wavelengths shorter than 506 nm, thus giving the possibility to divide the optical path into two different wavelength groups. The distance the two new beam components had to travel was ensured to be equal. Finally, the two beams were recombined by an identical dichroic mirror as the one splitting them and focused on the sCMOS chip side-by-side by the second achromatic doublet. The choice of achromatic doublets was made to reduce chromatic aberrations and focus different wavelength rays at the same point. It is noted that additional emission filters (e.g. Semrock, FF01-479/40 and FF01-542/27 for FRET experiments), were introduced in each path to narrow the transmitted wavelengths around the peak value of the desired emission spectra of the fluorophores used in experiments. Furthermore, the two dichroic mirrors were mounted on flip supports to easily switch to a single image detection scheme, also thanks to the possibility to enlarge the slit through a micrometer screw. Figure 3 shows a detailed schematic of the fluorescence microscopy part of the apparatus as well as the simultaneous acquisition of the “acceptor” and “donor” image channels in a sample containing Human Embryonic Kidney (HEK) cells which were transfected with an actin FRET force sensor (cpstFRET).

### 2.2 Optical Manipulation

An optical tweezers system was integrated to the setup (Fig. 4) to precisely apply external forces on cell membranes by trapping microscale beads and bring them in contact with the cells. Optical manipulation was accomplished by a diode laser (Lumics LU808M250), emitting at 808 nm with FWHM of 0.5 nm and maximum power *P*_*max*_ = 250 mW. This choice of wavelength was to minimize photodamage effects in living cells which are significantly reduced in the near-infrared region where the absorption coefficient is of the order of 10−4 [30], [31]. In experiments involving cells in vivo, it is essential to prevent the increase of the reactive oxygen species (ROS) level. Even though these molecules are fundamental for cell homeostasis and signalling, an increase in their levels originating from the interaction between light and the aqueous environment of a cell, can dramatically damage the cell structure. After the writing of this paper, the laser was exchanged with a laser (Thorlabs BL976-PAG900) of comparable cost and quality, slightly longer wavelength (976 nm), but considerably higher power (900 mW). The diode laser was driven by a compact laser diode driver (Thorlabs CLD1015). The trapping beam was directed to the microscope by being reflected by a piezo-driven steerable mirror. The piezoelectric mirror mount (PI N-480 PiezoMike) had an angular range of ϑ = ±5° (± 87 mrad) and minimum angular resolution of 0.3 μrad. This was conjugated to the back-conjugate focal plane of the microscope objective through a *M* = 10X magnifying telescope. In this way, the beam size was adjusted to slightly overfill the back aperture of the objective and the rotational movement of the mirror translated to linear movements of the trap in the sample plane. The lateral movement of the trap *x* ~ ϑ ∙ *f*_*o*_⁄*M*, where *f*_*o*_ = 3.33 mm is the focal length of the objective [11], could then span from a minimum lateral displacement of 0.1 nm to a maximum displacement of ±28 μm. Finally, the trap beam was directed to the microscope objective by a dichroic mirror (Chroma ZT775sp-2p). The Nikon Eclipse Ti-U microscope stative allowed the installation of an extension turret in between the fluorescence dichroic wheel and the microscope objective, which conveniently accommodated the tweezers’ dichroic mirror. Measurements were taken to produce a calibration curve describing the relation between the diode current and optical power of the beam, before and after the microscope objective. The power measured after the objective set the power incident on the sample and, thus, the stiffness of the trap. Typically, the power used in living cell experiments was 175 mW at the source output or 20 mW incident at the sample by setting a diode current at 250 mA.

**Figure 4:**
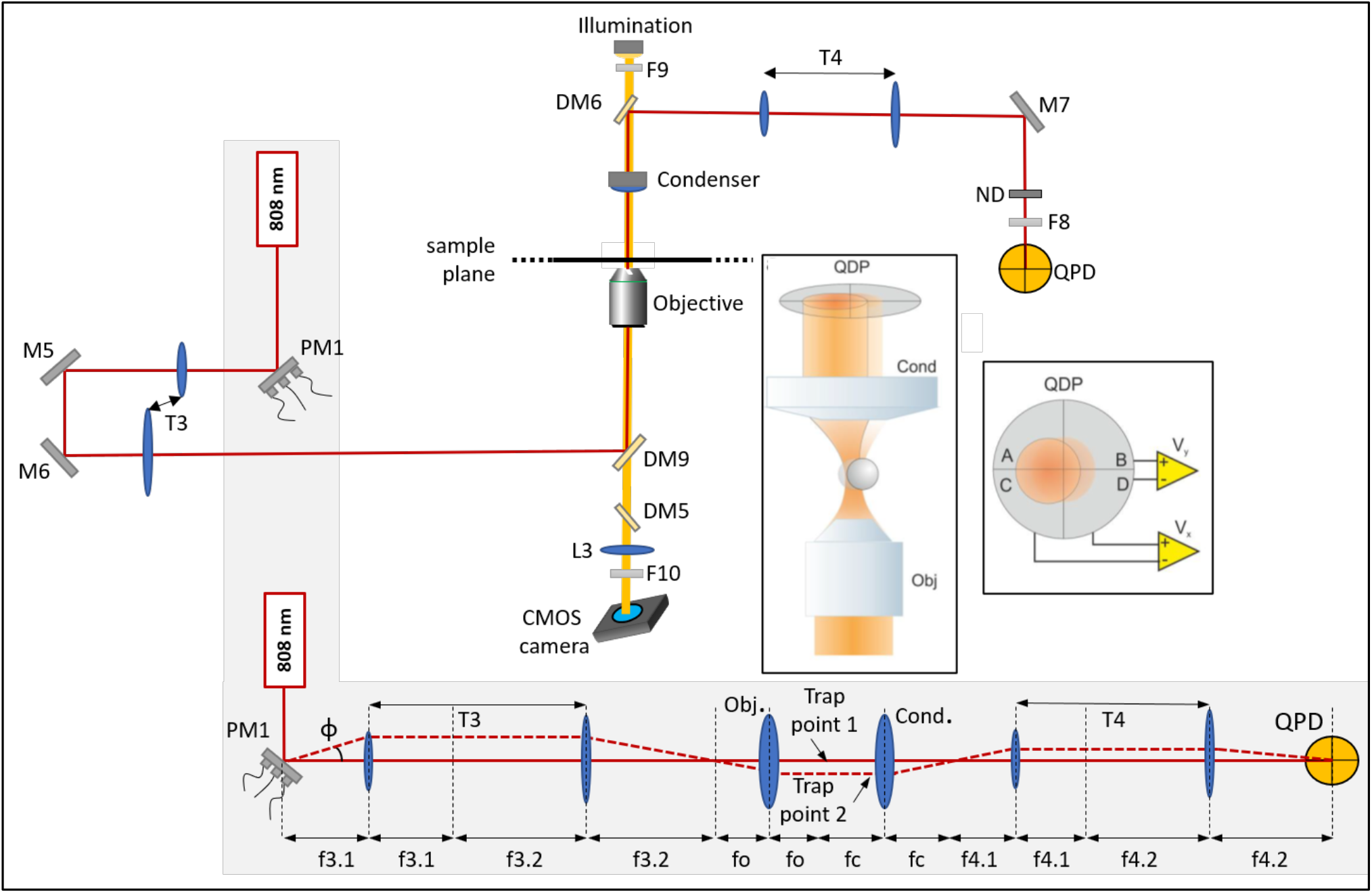
Schematic of the optical manipulation part of the experimental apparatus. An 808 nm laser source was used for the optical tweezers system. The piezo-electric mirror PM1 was used to stir the trapping beam to move the optical tweezers in the sample plane. The bottom grey area shows how the rotation of the piezo-electric mount and the placement of the optics can result to a new trap position. f3.1 and f3.2 denote the focal lengths of the lenses consisting the telescope T3, fo is the focal length of the microscope objective, fc of the condenser and f4.1 and f4.2 the focal lengths of the lenses creating telescope system T4. The beam was past through the telescope system T3 to enlarge the beam to overfill the microscope objective back aperture. A dichroic mirror DM9 directed the trap beam to the microscope objective. After passing through the sample, the beam light was collected by the condenser and reflected by the dichroic mirror DM6 towards the QPD. The beam was first passed through telescope T4, an ND filter and notch filter F8 before finally reaching the QPD. Inset: Interaction of scattered and unscattered light creates an interference pattern depending on the relative position of the trapped sphere with respect to the trap. The intensity distribution can be measured by means of the differential voltage signals by the QPD.

A quadrupole detector (QPD) (OSI Optoelectronics SPOT-15-YAG) with custom-built 30 kHz bandwidth electronics was used to detect and record the position of the trapped beads by collecting the interference pattern formed at the condenser back-focal plane[11]. The light was directed to the QPD by means of a dichroic mirror (Chroma ZT775sp-2p) reflecting wavelengths between 775 – 1200 nm mounted on top of the condenser lens. We used a condenser lens with a relatively low numerical aperture (NA = 0.52), to allow the use of cell culture dishes and a microscope incubator for live cell imaging (Okolab H301 chamber with UNO-T-H-CO2 all-in-one controller). The detector was conjugated to the back focal plane of the condenser through a demagnifying telescope (T4) to reduce the beam dimension to the detector active area and maintain detector alignment when the beam was steered by the piezo mirror (see Fig. 4, bottom). The later position of the trapped objects was obtained from the differential signals Vx and Vy, while the axial position from the total intensity [32].

Calibration of the optical tweezers system was performed by analysing the trapped bead position frequency spectra [33]. This was essential to quantify forces and displacements experienced by living cells. The calibration allowed to determine both the trap stiffness *k* and voltage-to-position photodetector conversion factor *β* [33]. Determining the conversion coefficient is equivalent to measuring the sensitivity of the photoelectric detector: *α* = 1/*β*. Since *β* = *x*/*V*, the power spectrum of the output voltage *V*(*t*) of the photoelectric detector can be obtained from the power spectrum of the position *x* of the trapped bead. Using the following formula:

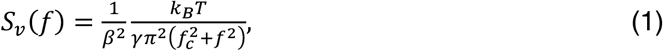

where *k*_*B*_ is the Boltzmann constant, *T* is the temperature, *γ* = 6*πηR* is the viscous drag coefficient, for a sphere of radius *R* immersed in a fluid of viscosity *η* and *f*_*c*_ is cut-off frequency, trap calibration can be obtained by fitting the voltage power spectra recorded while a particle is stably trapped by the optical tweezers with the above equation, using the least squares method. An example of frequency spectra obtained for a trapped 1 μm polystyrene bead suspended in water at room temperature is shown in Fig.5. The cut-off frequency *f*_*c*_ and the conversion factor *β* are the parameters that are chosen as free parameters in the fitting procedure. The obtained *f*_*c*_ value can be then used to obtain the trap stiffness *k* by:

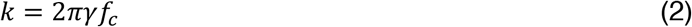

**Figure 5:**
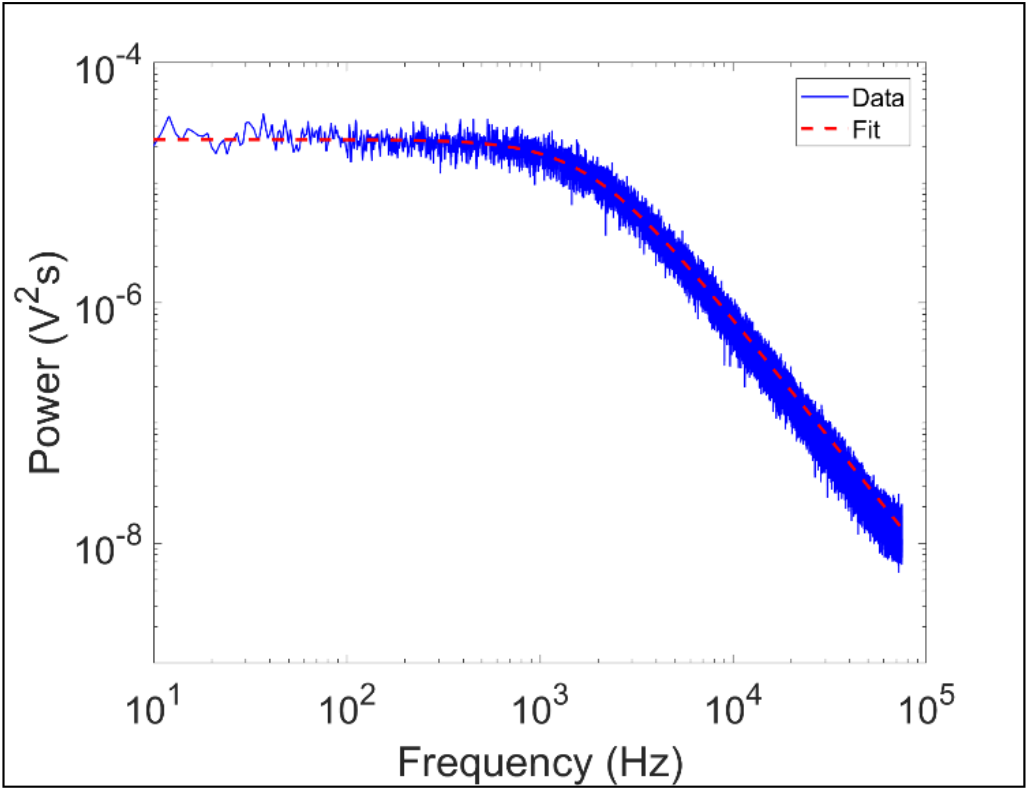
Example of a power spectrum (blue) for a 1 μm diameter polystyrene sphere immersed in aqueous solution at room temperature and trap laser power 28 mW. The red dashed line represents the fit function used for the optical tweezers calibration The trap stiffness *k* and the conversion factor *β* in this case were found to be 0.1 pN/nm and 24.0 nm/V in this case.

We tested the linearity of the trap stiffness and the independence of the voltage-to-position conversion factor with the laser power, both for polystyrene and silica beads.

## 3 Characterization of the experimental setup

### 3.1 Single Molecule Detection

Fluorescence excitation of organic chromophores and fluorescent proteins was obtained using total internal reflection fluorescence (TIRF) to follow processes occurring near the cell membrane, or with an optimized highly-inclined light optical (HILO) sheet to image intracellular or nuclear events with high signal-to-noise ratio [34]. In order to follow mechanotransduction signal at the molecular scale, the sensitivity of the apparatus owned to be able to detect single chromophores and localize them with few nanometre resolution. To establish the resolution of the apparatus, measurements based on single chromophore localization were performed aiming to accurately extract the coordinates of each dye in the sample plane. The localization of single fluorescent molecules was determined through fitting of the point spread function (PSF) with a bidimensional Gaussian [35], [36] centered at (*x_0_, y_0_*):

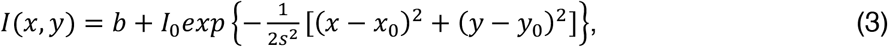

where *b* is the background intensity, *s* is the two-dimensional standard deviation, and *I*_*0*_ is the peak intensity.

The localization uncertainty can be reduced by the increase of collected photons and by minimizing noise factors [35]. The nanometre localization capability of the setup was demonstrated by using Alexa Fluor 568 and Alexa Fluor 647, with absorption and emission maxima at 578 nm and 603 nm, and at 650 nm and 665 nm, respectively. All samples were incubated in specific flow chambers (Fig. 6a). First, a microscope slide and a 18 mm × 18 mm glass coverslip with thickness of 170 μm were cleaned with pure ethanol and then dried with under nitrogen flow. Two flow chambers were then created by adding three stripes of double-stick tape to the slide surface and attaching the coverslip to the tape. Each chamber measured ~ 15 μL of volume. 20 μL of Biotin-conjugated Bovine Serum Albumin (bio-BSA) 1 mg/ml was flowed in each chamber and incubated for 5 minutes. Next, the chambers were washed with 60 μL of Phosphate buffer (PB) 50 mM. Subsequently, 20 μL of the two streptavidin-conjugated alexa-fluor dyes were separetely flowed in the two chambers and incubated for 3 minutes. The concentrations chosen for Strepatavidin-Alexa Fluor 647 was 0.2 ng/mL and for Strepatavidin-Alexa Fluor 568, 0.7 ng/mL to obtain separated single emitters attached on the coverslip surface. Finally, the chambers were washed with 100 μL of imaging buffer (928 μL PB, 10 μL DTT 1M, 40 μL Glucose Oxidase 5 mg/ml, 10 μL Catalase enzyme 5 mg/ml, 12 μL Glucose 250 ng/ml). Careful wash is essential in preventing the presence of background fluorescent molecules during the imaging process, while the oxygen scavenger enzymes in the imaging buffer assure reduced oxygen levels in the sample to minimize blinking and photobleaching phenomena.

**Figure 6:**
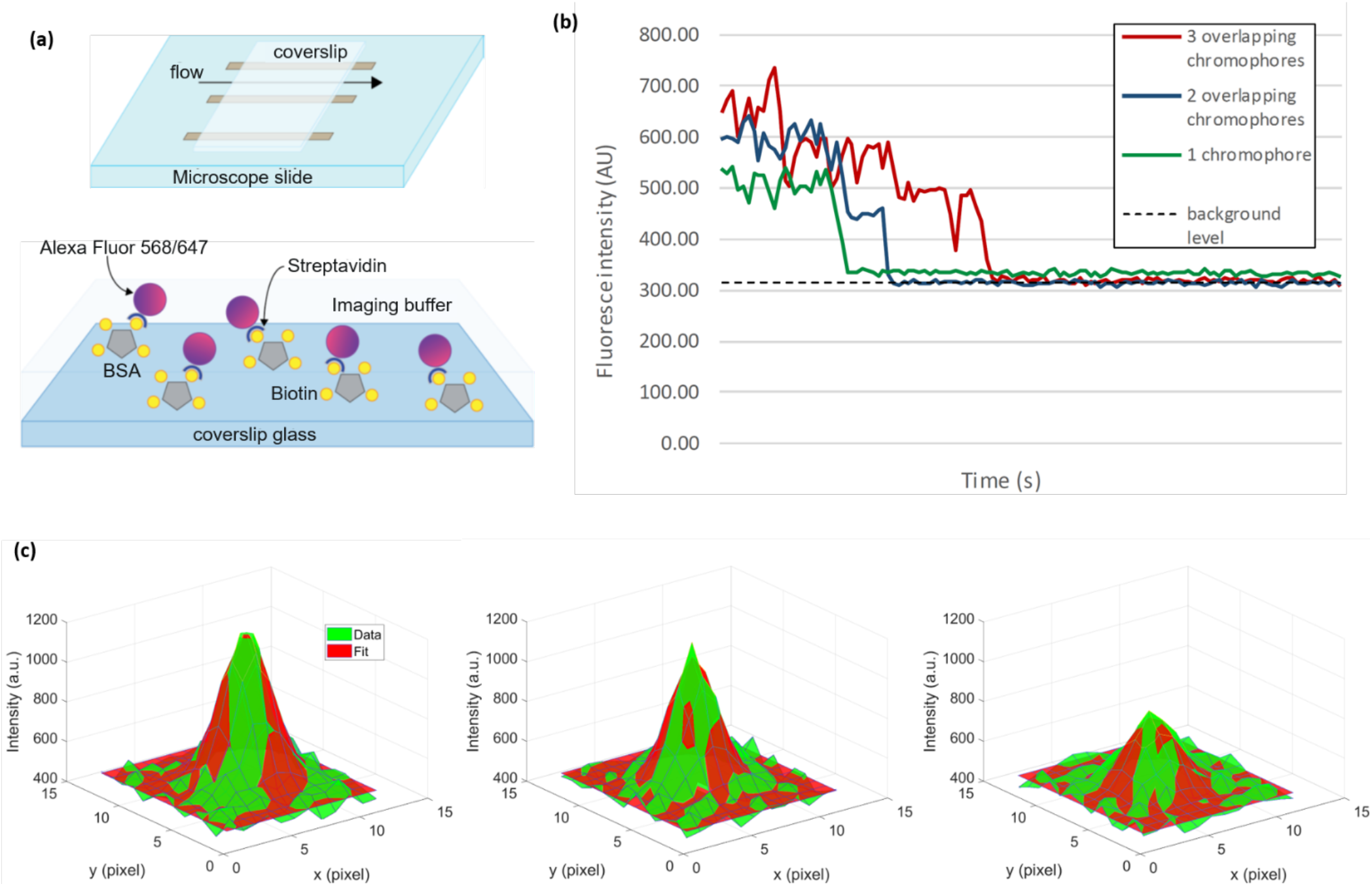
**(a)** Top: Flow chamber. Bottom: streptavidin-Alexa Fluor molecules were specifically bound to the coverslip through biotinylated BSA. **(b)** Emission intensity dropping in step-like fashion implying that the source of emission consisted of a cluster of multiple dyes, each one photobleaching in time. **(c)** Time-evolution of FIONA algorithm tracking of single chromophores. The green colour represents the recorded data while red is the gaussian fit providing the centre of mass coordinates. The intensity drops observed between frames is due to the photobleaching of one dye.

The sCMOS camera was set to acquire 200 consecutive frames with 500 ms integration time. The acquisition of data was carried out for the three different illumination configurations: wide field, HILO and TIRF. The chromophores were illuminated with excitation intensity at *I* = 100 W/cm^2^. HILO and TIRF are considered as high signal-to-noise ratio fluorescence microscopy techniques due to the reduction of background noise rising from out-of-focus fluorescence. These illumination schemes provide higher excitation intensities compared to wide-field microscopy. Optimized TIRF microscopy, i.e. when the incident angle *θ*_*i*_ equals the critical angle for total internal reflection, results to *I*_*TIRF*_ = 4*I* = 400 W/cm^2^ which was comparable to intensities produced by optimized HILO microscopy.

In order to ensure that the localization analysis was conducted on single chromophores the temporal evolution of fluorescence emission was studied. In some cases, emission intensity dropped in a step-like fashion with time, thus implying that the source of emission was a group of agglomerated dyes instead of a single molecule (Fig 6b). Each step, in decreasing intensity denoted the photobleaching of a chromophore belonging to these groups. Furthermore, in these situations quenching and blinking effects originating from the interaction between the different chromophores also resulted in irregular intensity readings. Parts of the collected data that contained such dye groups were not considered for localization analysis. Moreover, the absolute position of each dye had to be corrected to account for thermal drifts. Therefore, we used as a reference the coordinates (*x_ref_*, *y_ref_*) of a single chromophore in the field of view which emitted at constant average intensity during all frames and calculated the position of the other chromophores as the relative position (*x*^*r*^ = *x* − *x*_*ref*_, *y*^*r*^ = *y* − *y*_*ref*_).

We then evaluated the localization accuracy for each chromophore as the standard deviation of its relative position and the mean localization accuracy for all the Alexa Fluor 647 chromophores analysed was evaluated to be 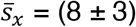 nm and 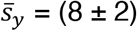 nm for wide field microscopy, 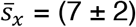 nm and 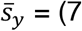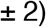 nm for HILO and 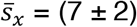 nm and 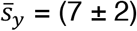 nm for TIRF. For Alexa Fluor 568 were found 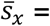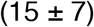 nm and 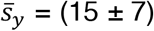 nm, 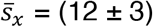 nm and 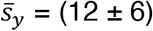 nm, and 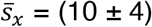 nm and 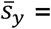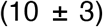 nm for the different illumination schemes, respectively. The errors given here are standard deviations from the average value.

### 3.2 Cell Membrane Stimulation

In mechanotransduction experiments, optical tweezers can be used to manipulate specific receptors on live cells’ membrane by applying force through a trapped bead. Consequently, this can induce structural modifications in the membrane receptor and trigger intercellular processes. Fig. 7a shows an illustrative cell manipulation experiment in which a fibronectin coated polystyrene bead is attached to the cell membrane and being pulled away from it. A cell membrane tether is clearly visible, connecting the bead to the cell while a force is applied [13]. The QPD signal was recorded through this process and the applied force on the membrane during pulling the tether was calculated (Fig. 7b). The trapped bead displacement was determined by *β* obtained from fitting Eqn. 1 to the power spectra, and the difference in voltage signal as recorded by the QPD for each step i.e. *x_i_ = β·ΔV_i_*. The force was then calculated for each step by using the trap stiffness as also obtained by the power spectra fitting. Finally, the mean force and displacement were calculated for all steps revealing a linear relationship between the two. The tether stiffness could be calculated with an error by a linear fit (Fig. 7c)

**Figure 7:**
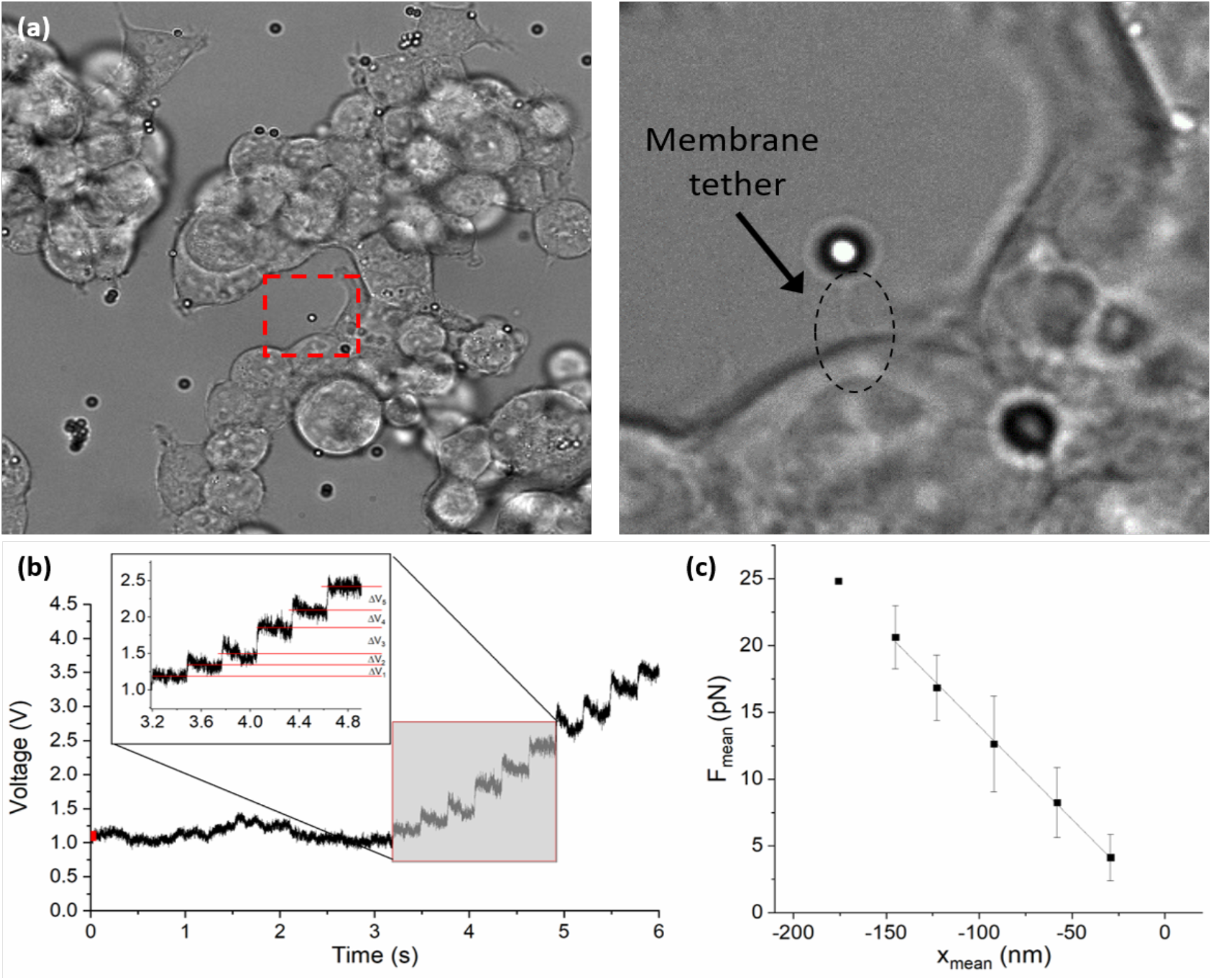
**(a)** Fibronectin coated bead comes to contact with the cell membrane via the optical tweezers. The bead is then pulled away from the cell creating a membrane tether while applying force to the membrane. **(b)** The QPD signal as recorded during step-wise motion of the trap, beam power 28 mW. The bead follows the movement of the trap with some delay. **(c)** Mean force vs mean bead displacement revealing a linear response. Tether stiffness *kteth.*was found to be 0.139 ± 0.001 pN/nm in this case.

Cell manipulation experiments can be then combined with the imaging of intracellular signals by exploiting the single molecule sensitivity of our setup. Examples include the imaging of mechanically-triggered intracellular Ca+2 fluxes [37], mechanically-induced changes in protein expression using a fluorescent reporter[24] or changes in mRNA expression that can be visualized with single molecule sensitivity using specific reporter such as the MS2 system [26][27]. As reported in the next paragraph, we can also exploit the combined fluorescence imaging to image mechanical signals and changes in intracellular tension by using FRET-based force sensors inserted in specific cytosolic proteins.

### 3.3 Force imaging through FRET-based genetically encoded force sensors

#### 3.3.1 FRET Calibration using FRET standards

The phenomenon of Fluorescence Resonance Energy Transfer (FRET) can be used to study protein-protein or protein-cell membrane interactions inside living cells. There is a relevant number of different methods to measure FRET using a multitude of FRET indexes that relates to the actual FRET efficiency. Since the FRET efficiency is usually undefined or subject to relevant systematic errors, it is very difficult to compare FRET measurements obtained with different methods and setups. A possible solution to this problem comes from the development of genetic constructs encoding fluorescent proteins (FPs) pairs with known FRET efficiency, which can be used as “standards” to validate and calibrate FRET in different imaging systems with high reproducibility [38].

Koushik et al. used a Cerulean/Venus donor/acceptor pair to engineer three FRET standards with growing distance between the donor/acceptor pair: C5V, C17V and C32V [38]. C stands for Cerulean (donor, *λ*_*exc*_ = 433 *nm*; λ_25_ = 475 *nm*) and V stands for Venus (acceptor, *λ*_*exc*_ = 515 *nm*; λ_25_ = 528 *nm*), while the number indicates how many amino acids are placed in the linker between the FPs. We used these FRET standards to calibrate our FRET imaging system and relate our measurements of FRET index to FRET efficiency values obtained by Koushik et al [38].

In the measurements presented here, C5V, C17V and C32V were transfected inside HEK cells, which were allowed to adhere on a glass coverslip. Images were taken sequentially: first, by exciting the sample with the 445 nm laser with a 50 ms exposure time to acquire the FRET and Donor Channels; then, by illuminating with the 514 nm laser with 50 ms exposure time to image the Acceptor Channel. A custom-made MATLAB script was prepared to readjust the images and correct for chromatic aberrations. The script allows to select the region of interest in the donor channel on which the user wants to perform the correlation process. This is done through a specific MATLAB algorithm which performs a 2-D cross-correlation between two images, *A* (in the donor channel) and *B* (in the acceptor channel). This was done to find the exact point where a section of the first image is superimposed onto the same section in the second image, in order to find the regions where the two signals are identical. The script also scales the images to readjust the dimensions of each channel by using the brightfield image as reference. Finally, as a result, the script automatically builds an image stack composed by the three channels aligned (Fig. 8). After cross-correlation and scaling, all images were analysed using the ImageJ PixFRET plug-in [39]. This plugin returns a final image where a FRET index is calculated and displayed in each pixel with a colour map (Fig. 8). The FRET index is then normalized to find a ratiometric FRET index which is independent from the density of the FPs and is given by the formula:

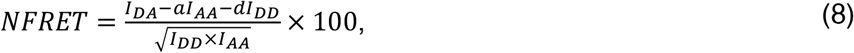

where *I*_*DA*_ is the fluorescence intensity obtained through excitation of the donor and measured in the acceptor channel, *I_AA_* is the intensity measured in the acceptor channel when excitation wavelength of the acceptor is used, and I_DD_ is the intensity measured in the donor channel upon donor excitation. The parameters *a* and *d* are the spectral bleed-through ratios, which represent the portion of fluorescence emission collected in the FRET channel which is not produced by the energy transfer. In particular *a* is the emission of the donor in the acceptor channel, while *d* is the excitation of the acceptor using the donor excitation wavelength. The final FRET index of each cell is found by using the “Analyse Particles” function in ImageJ which returns the particle (cell) area, the mean value of FRET index of the cell, its standard deviation and standard error. The FRET index of each standard construct is then calculated simply as the mean FRET index value of all the cells of that specific construct, with the associated standard error.

**Figure 8:**
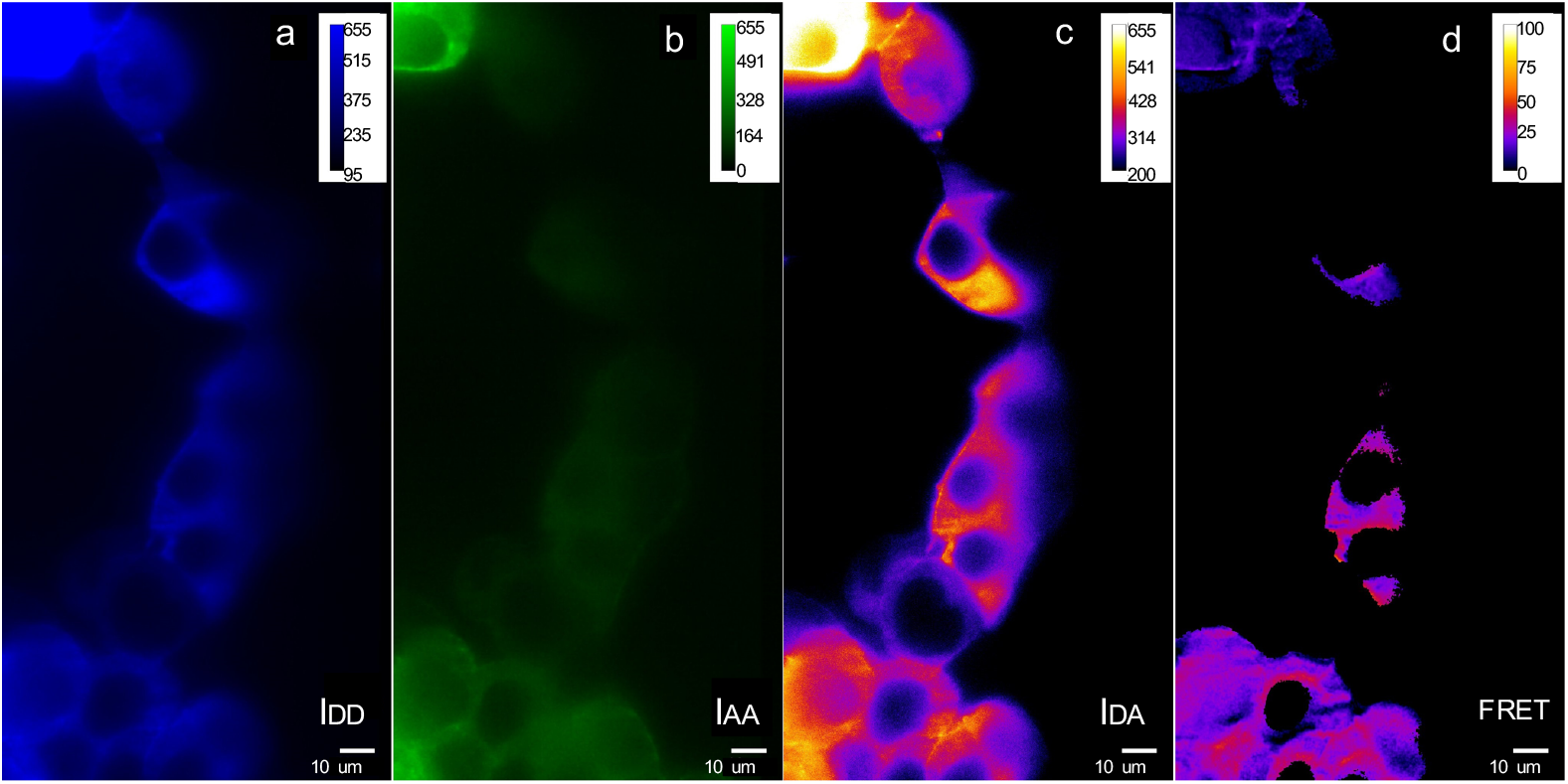
Images of FRET standards and FRET index. The image shows HEK cells transfected with the C32V FRET standards. **(a)** The donor channel (excitation with the 445 *nm* laser and emission in the range 460 *nm* < *λ*_*em*_ < 500 *nm*). **(b)** The acceptor channel (excitation with the 514 *nm* laser and emission in the range 530 *nm* < *λ*_*em*_ < 560 *nm*). **(c)** The FRET channel (excitation with the 445 *nm* laser and emission in the range of the acceptor channel). **(d)** Ratio metric FRET index image calculated through equation (8). The power of both lasers on the sample plane was 2.5 mW, 50 ms integration time.

FRET indexes measured as described above are shown in the histogram in Figure 9a. The linear trend observed in both our measurements of FRET index and the FRET efficiencies obtained in [38] (Fig. 9b) within the measured range, allowed for a calibration curve relating the measured FRET index with the calibrated FRET efficiency, for a better comparison with experiments performed with different methods (Fig. 9c).

**Figure 9:**
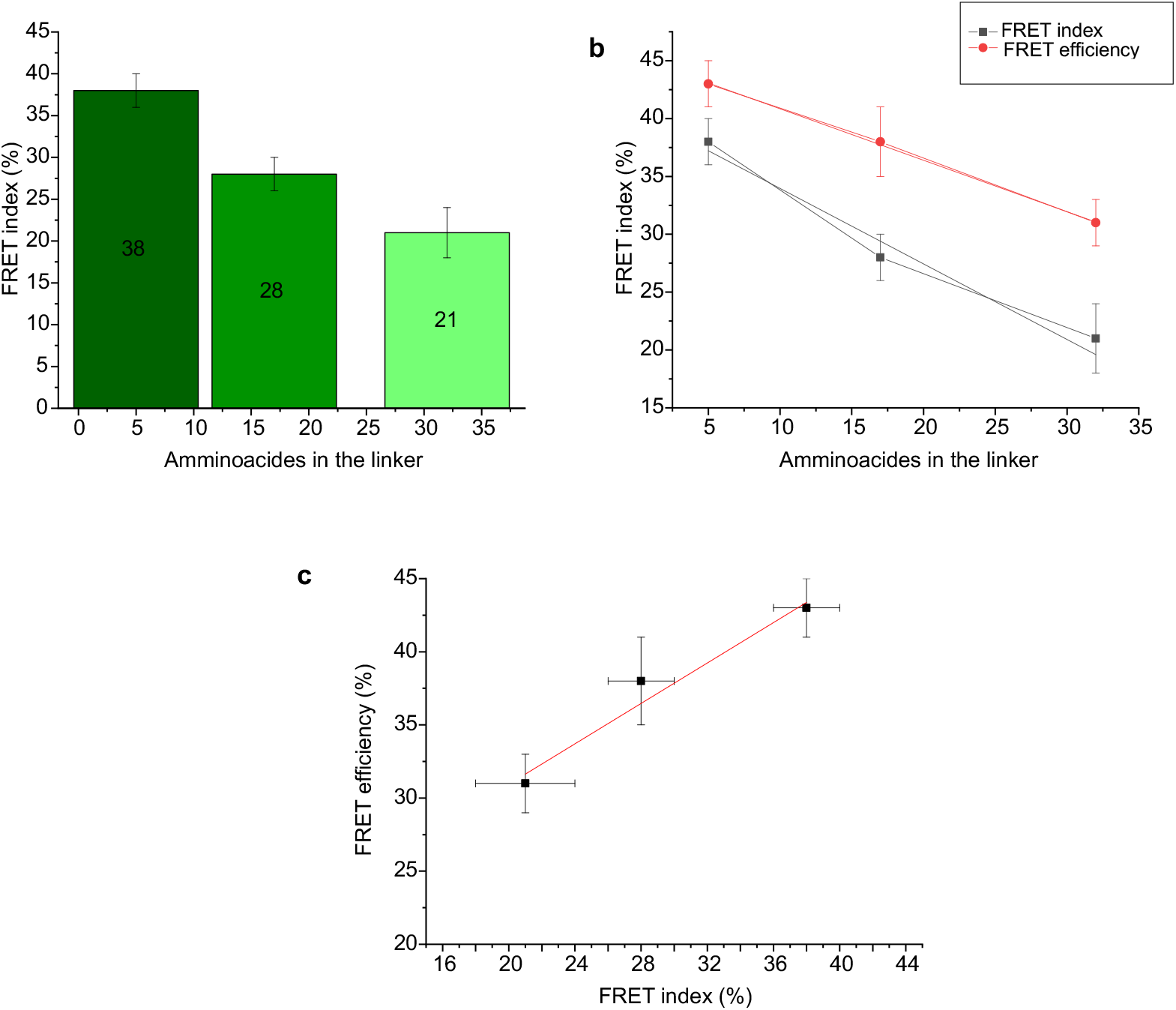
Characterization of FRET standards. **(a)** FRET indexes of the C5V, C17V and C32V standards decrease with increasing distance between the fluorophores of the FRET pair. **(b)** FRET index of C5V, C17V and C32V vs calibrated FRET efficiency (N=10 cells per construct). The linear fit of FRET index gives an adjusted R- square value of 0,933. **(c)** Calibration curve to relate FRET index measurements with the calibrated FRET efficiency. The linear fit between FRET efficiency and FRET index gives a slope of (17 ± 6) and an intercept of (0,7 ± 0,2), with a Pearson’s coefficient value of 0,98.

#### 3.3.2 Measuring tension on the actin cytoskeleton

After calibration, a study of a specific sensor inserted into F-actin was conducted [17]. The sensor used in these experiments was based on the donor-acceptor relative orientation, unlike the FRET standards used previously which were modulated by the distance between the two. This sensor is called cpstFRET (circularly permuted stress sensitive FRET) and it was developed based on the existing stFRET sensor [18]. cpstFRET is composed of five amino acids that link two circularly permuted FPs that are parallel at zero force and are supposed to twist towards a perpendicular configuration under tension, decreasing FRET efficiency values. As a consequence, this sensor is much smaller than the ones based on linear springs [16].

The actin probe used, named actin-cpstFRET-actin (AcpA), is shown in Fig. 10a and it is constituted by the dipole orientation-based sensor cpstFRET flanked by two β-actin monomers, which label F-actin efficiently. A zero-force control sensor called cpstFRET-actin (cpA) was also used (Fig 10a). This sensor has only one actin monomer, leaving the other end of the probe free to move and not be subjected to forces [17]. Both sensors were successfully transfected into HEK cells which were grown on different substrates.

**Figure 10.**
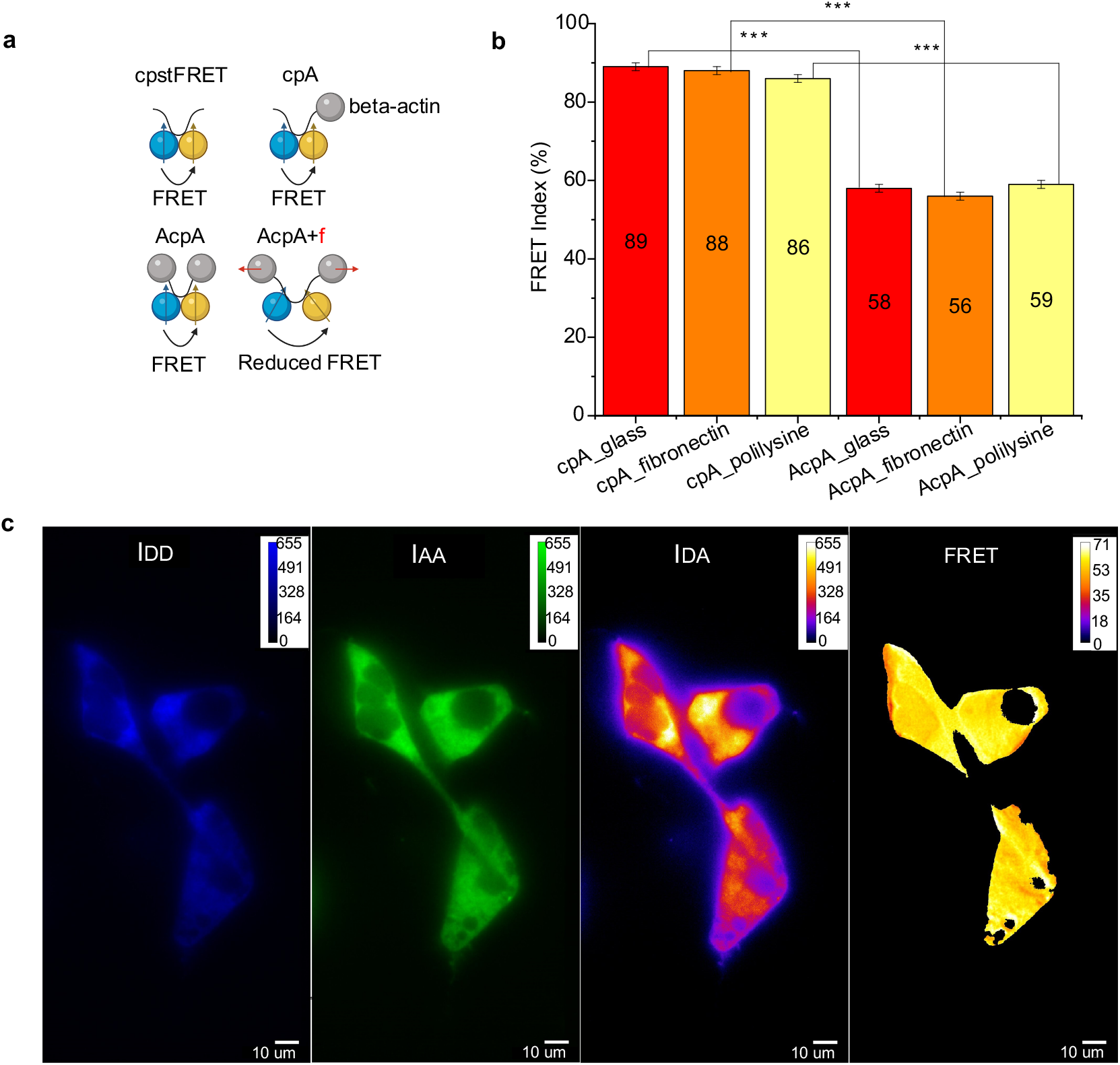
**(a)** Sketch of the cpstFRET sensor and how it is inserted into F-actin. “cpA” is a cpstFRET-β-actin construct, where one actin monomer is bound to the C-terminal of the cpstFRET sensor. “AcpA” stands for “Actin-cpstFRET-Actin” and is a construct where two actin monomers are linked by the cpstFRET sensor. “f” represents an external force, which induces rotation of the emission dipoles of the FPs, thus resulting in lower FRET [17]. **(b)** FRET indexes of cpA and AcpA in HEK cells grown on glass slides (red), glass slides covered with fibronectin (orange) and glass slides covered with polylysine (yellow). All the force-free cpA sensors have significantly higher indexes compared to the AcpA sensor. According to a two-sample t-Test, FRET index of AcpA on glass (*N* = 20) is not significatively different from FRET indexes of AcpA grown on fibronectin (*N* = 23, *p* = 0,43) and polylysine (*N* = 20, *p* = 0.79). **(c)** Example of a HEK cell transfected with the AcpA sensor. Images were taken in the three channels with an integration time of 50 ms and the power of both lasers at the sample plane was 8 mW. Last image of panel C is the FRET index image calculated by the PixFRET plugin as described above.

First, the FRET efficiencies of our sensors (both the sensor and the control) transfected into HEK cells and grown on bare microscope coverslips were evaluated. All measurements were taken while keeping all experimental parameters constant compared to the previous measurements on the FRET standards. Consequently, the data were analysed using the PixFRET plug-in, from which the mean values of normalized FRET indexes were obtained along with the calculated standard errors. As expected, a high FRET index of (89 ± 1)% for the force-free actin sensor (cpA) was obtained, while this value decreased to (58 ± 1)% for the AcpA sensor, which is subject to tension (Fig. 10b). Next, cells with actin sensors were grown on microscope glass slides covered with polylysine, which facilitates cell adhesion, and fibronectin, which stimulates focal adhesion (FA) formation and possibly stimulates higher mechanical tension on the actin cytoskeleton.

The results (Fig. 10b) revealed that there was no significant difference between the actin sensors grown on different substrates. A possible explanation derives from the fact that in these series of measurements, the microscope objective was not focused at the glass-cell interface where FAs form, but instead deeper into the cell. Thus, observations made originated from parts of the cytoskeleton that were possibly not significantly influenced by focal adhesions.

#### 3.3.2 Measuring tension on α-actinin

α-actinin is a cross-linking and bundling protein that constitutes the backbone of contractile actin bundles and stress-fibers. An α-actinin force sensor was also investigated. In this sensor, called Actinin-M-cpstFRET, the cpstFRET cassette is inserted between spectrin repeat domains 3 and 4 towards the middle of α-actinin. On the other hand, the zero-force control called Actinin-C-cpstFRET is created by binding the cpstFRET construct to the C-terminal of α-actinin. A scheme of these two sensors is shown in Fig. 11a [17].

**Figure 11.**
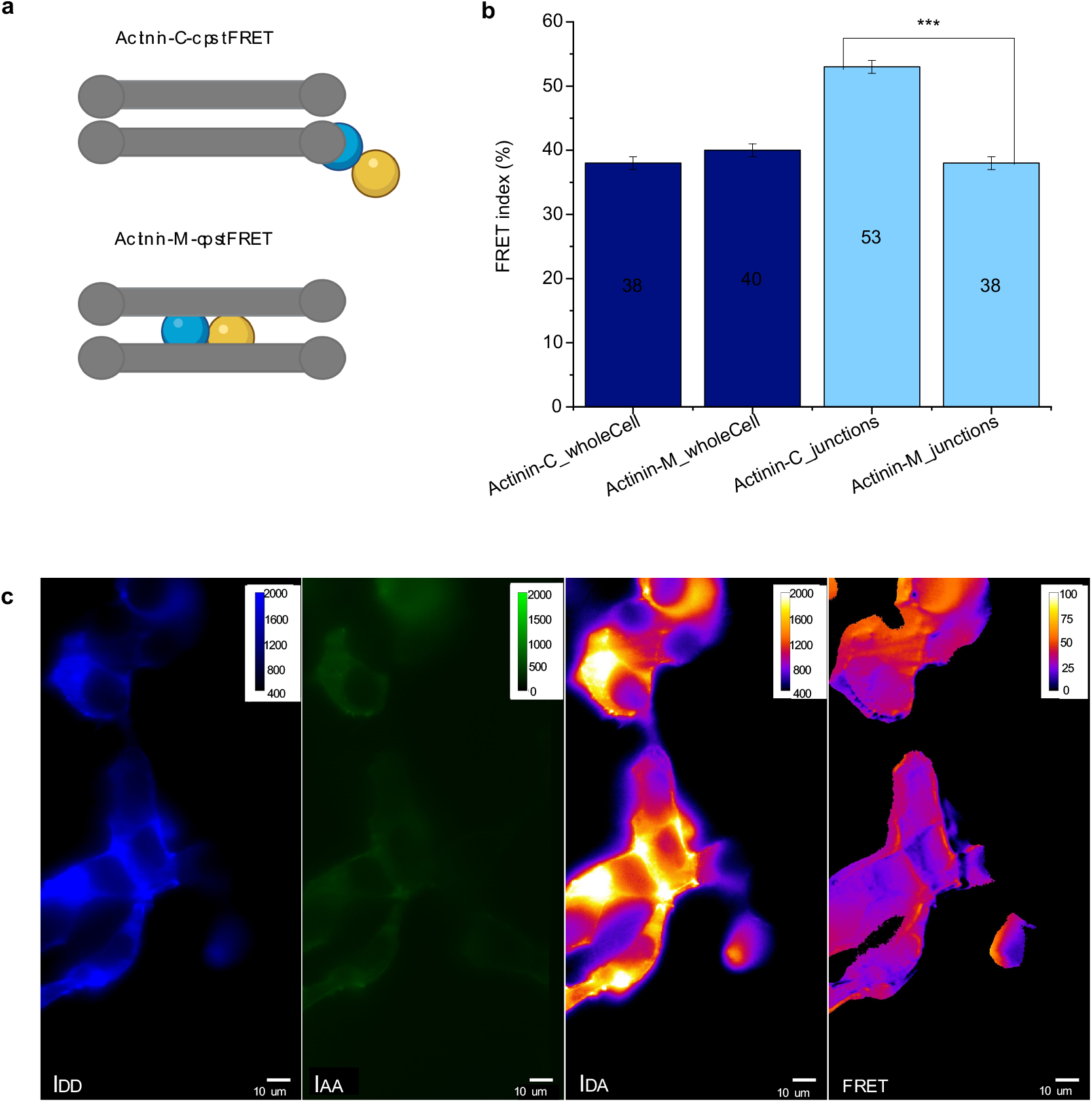
**(a)** Sketch of the two α-actinin sensors used. “Actinin-C-cpstFRET” is the zero-force control sensor, where cpstFRET is linked to the C-term of α-actinin. “Actinin-M-cpstFRET” is the force sensor, where cpstFRET is inserted in the middle of the α-actinin protein and is subjected to stress under tension [17]. **(b)** FRET indexes of Actinin-C-cpstFRET and Actinin-M-cpstFRET grown on microscope glass slides and measured through TIRF microscopy in the whole cell (blue, *N* = 123 for Actinin-C-cpstFRET, *N* = 140 for Actinin-M-cpstFRET) and only along cell-cell junctions (light blue, *N* = 131 for Actinin-C-cpstFRET, *N* = 103 for Actinin-M-cpstFRET). Analysis of cell-junctions only reveals a significant difference between the zero-force sensor and the sensor subjected to tension (two-sample t-Test, *p* < 0,05). **(c)** Image of Actinin-M-cpstFRET sensor in HEK cells acquired through TIRF illumination. Fluorescence emission is concentrated mainly along cell-cell junctions. Images were taken in the three channels with an integration time of 50 ms and the power of both lasers at the sample plane was 8 mW. Last image of panel C is the FRET index image calculated by the PixFRET plugin as described above.

Initially the FRET indexes of the sensors were characterised in HEK cells grown on microscope coverslips by TIRF microscopy. Measurements were taken with the same parameters as the previous experiments. The images obtained were analysed as usual with the PixFRET plugin. The results obtained are shown in Fig. 11b (blue). With 0.05 confidence level, there was no significant difference between the sensor under tension and the force-free control (*p* > 0.05 from the standard statistical two-sample t-Test). However, through a more careful inspection of the images acquired (Fig. 11c), it was found that the actin fluorescence was prevalently localized along cell-cell junctions in this case. For this reason, a new analysis was performed by selecting only the junctions’ area as the region of interest, where α-actinin was highly concentrated and probably under tension. As it can be seen by the results shown in Fig. 11b (light blue), the difference between the zero-force control and the sensor is significant (*p* < 0,05 with standard two-sample t-Test). As expected, the FRET index of the Actinin-M-cpstFRET sensor is lower than the Actinin-C-cpstFRET sensor, underlying that α-Actinin is affected by tension near cell-cell junctions.

## 4 Conclusions

The study of mechanotransduction in living cells involves the measurement of force in multiple cell locations with molecular specificity, as well as the simultaneous imaging of biochemical and genetic signals transduced by the cell. The complexity of these measurements requires specialized setups combining multiple techniques. Here we propose a combination of optical tweezers, FRET-based molecular tension microscopy, and fluorescence imaging with single molecule sensitivity to mechanically stimulate cells and simultaneously image the propagation of mechanical and biochemical signals inside the cell. We give details on the setup implementation to allow other researchers to reproduce the setup performance and we give protocols to test and calibrate the different components. The developments reported here will allow future studies on the transmission of mechanical forces from the outer cell membrane to the cell’s cytoskeleton and nucleus, and how those forces are transduced into other types of signals and cell responses, hopefully opening new possibilities for the understanding of mechanotransduction mechanisms in living systems.

## Acknowledgments

We thank Prof. Frederick Sachs for the kind gift of the plasmids and cells containing the actin and alpha-actinin force sensors and for the scientific discussion. This work was supported by the European Union’s Horizon 2020 research and innovation program under grant agreement no 871124 Laserlab-Europe, by the Italian Ministry of University and Research (FIRB “Futuro in Ricerca” 2013 grant n. RBFR13V4M2), and by Ente Cassa di Risparmio di Firenze.

